## Supplemental for "Structure of a HIV-1 IN-Allosteric Inhibitor Complex at 2.93 Å Resolution: Routes to Inhibitor Optimization"

#### Supplemental Figure Legends

**Supplemental Figure 1. ALLINI-induced polymers of HIV-1 IN $CCD^{F185K}$•$CTD^{220\text{-}288}$.** A. Shown are 100 µM mixtures of $CCD^{F185K}$•$CTD^{220\text{-}288}$ in the absence or presence of 100 µM BI-224436. In the presence of ALLINI, modest turbidity is observed. B. Shown is autocorrelation data from dynamic light scattering (DLS) for 1 µM BI-224436 alone (orange), 1 µM $CCD^{F185K}$•$CTD^{220\text{-}288}$ with DMSO (yellow), and 1 µM $CCD^{F185K}$•$CTD^{220\text{-}288}$•BI-224436 (purple). Strong scattering data arises when both protein domains and drug are present, coinciding with observed turbidity. C. Particle distribution analysis of autocorrelation data shown in panel B. Data were well-described by a unimodal model with a particle diameter near ~$10^4$ nm. D-E. Sedimentation velocity analytical ultracentrifugation (SV-AUC) analysis on the interaction of $CCD^{F185K}$ and $CTD^{L242A}$ (light blue) in the presence of BI-224436 (light blue). In the leftmost panels, fits of the experimental data (circles) to the Lamm equation are shown as lines; in middle panel the residuals from this fitting are shown. Every third boundary and are shown for clarity. Measurements were performed at 10-30 µM monomer concentrations at 20ºC. D&E**.** c(S) (D), van Holde-Weischet (inset, D), and c(M) (E) distributions were derived from the fitting of the Lamm equation to the experimental data collected, as implemented in the program SEDFIT, with an overall RMSD of <0.005 for all fits. $CTD^{L242A}$ data is shown in blue, $CCD^{F185K}$ in light blue, and complex data in purple. This analysis shows evidence of ternary complex formation in the presence of BI-224436.

**Supplemental Figure 2.** A. Packing of the crystallographic asymmetric unit, showing four ternary complexes, which in turn provides eight independent observations of the ALLINI binding site.

Representative sigma-weighted $2F_o$-$F_c$ electron density, weighted at 1σ, is shown. B. Superposition of the Cα-traces of apo CCD$^{F185K}$ (salmon) with CCD$^{F185K}$•BI-224436 (blue). Highlighted is the sidechain of Trp-131. The two chains with an RMSD of 0.3 Å over matching atoms, with only slight discrepancies in backbone atoms observed across the α4 region (residues 138-155). C. Superposition of the Cα-traces of apo CCD$^{F185K}$ (salmon), CCD$^{F185K}$•BI-224436 (blue), and CCD$^{F185K}$•BI-224436•CTD ternary complex (tan). Boxed in grey dotted lines and shown is inset is the α3 helice at the ALLINI binding interface, highlight the conformational change occurring with Trp-131 upon complex formation. D. Comparison of the Trp-131•Arg-224 cation-π interaction at ALLINI sites 1 and 2. On the left, a superposition of the Ca traces for the four observations of the first ALLINI site is shown, with preservation of this interaction observed across all four protomers. On the right, a similar rendering of the Cα traces for the four observations of the second ALLINI site. In this second site, significant variation and perturbation of the cation-π interaction is observed.

**Supplemental Figure 3. Sequence conservation of the CCD and CTD domains at the ALLINI binding interface.** The DNA sequence alignments of HIV Integrase were extracted from the 2019 HIV Sequence Alignments published at the Los Alamos database[1]. A non-redundant collection of 4326 sequences representing all strains, as well as 2471 sequences of Group M from HIV-1/SIVcpz organism were extracted and translated. The HIV IN sequences of subtypes B and C, on the other hand, were selected from the collection of Group M sequences. Protein sequences were aligned against HIV-1NL4-3 (GenBank: AAK08484.2) using standalone Clustal Omega (version 1.2.3)[115-117]. Conservation scores of sequence alignments were calculated using the entropy-based method from AL2CO[111] program implemented in the UCSF ChimeraX[112]. Regions of high conservations are shown in magenta while regions of low conservations are in cyan. Top: comparison of sequence conservation of the CCD and CTD domains at the ALLINI binding interface between HIV-1 all strains, Group M without circulating recombinant forms (CRFs), Subtype B, and Subtype C. Consistent among the panel, is the high sequence conservation observed along the CTD, which includes Trp-235, Lys-266, and Tyr-226. Glu-170, Ala-169, and Thr-174 at the CCD are moderately conserved. Bottom: comparison of sequence conservation of the CCD dimeric interface at the ALLINI binding site. Here, the region of low conservation, which includes Gln-95, Thr-125, and Thr-124, remained consistent across the board.

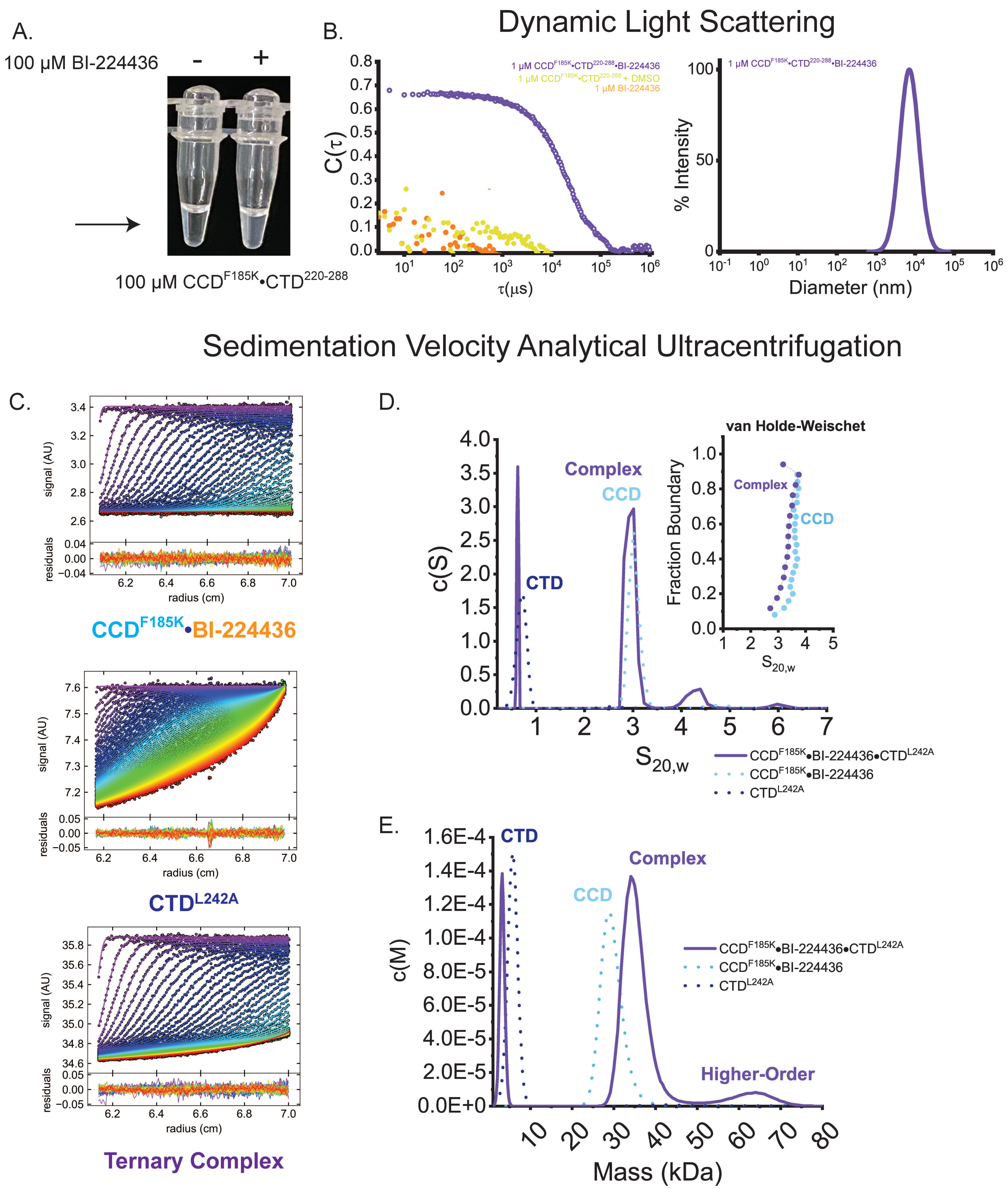


Supplemental Figure 1.

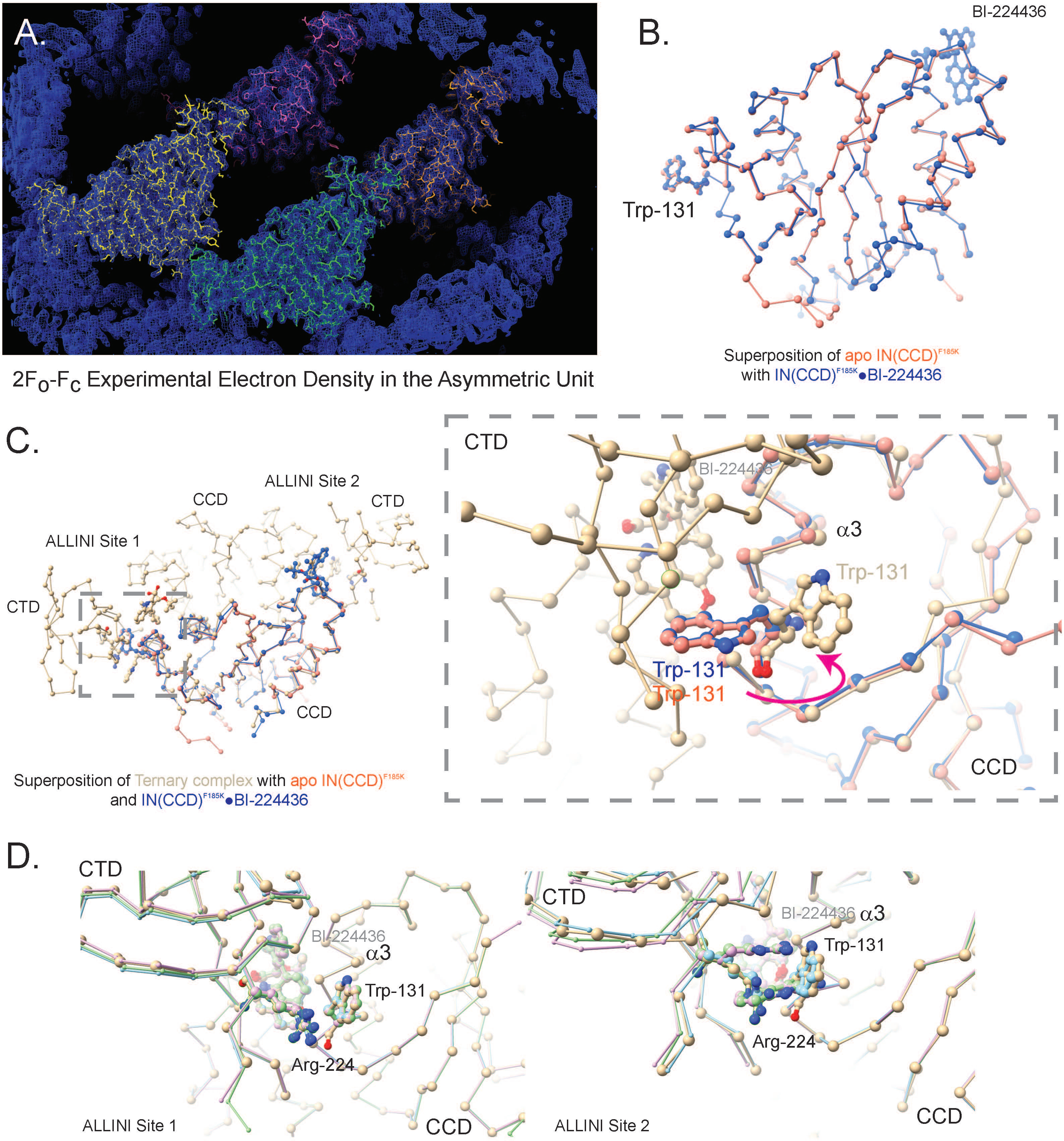


### Suppemental Figure 2.

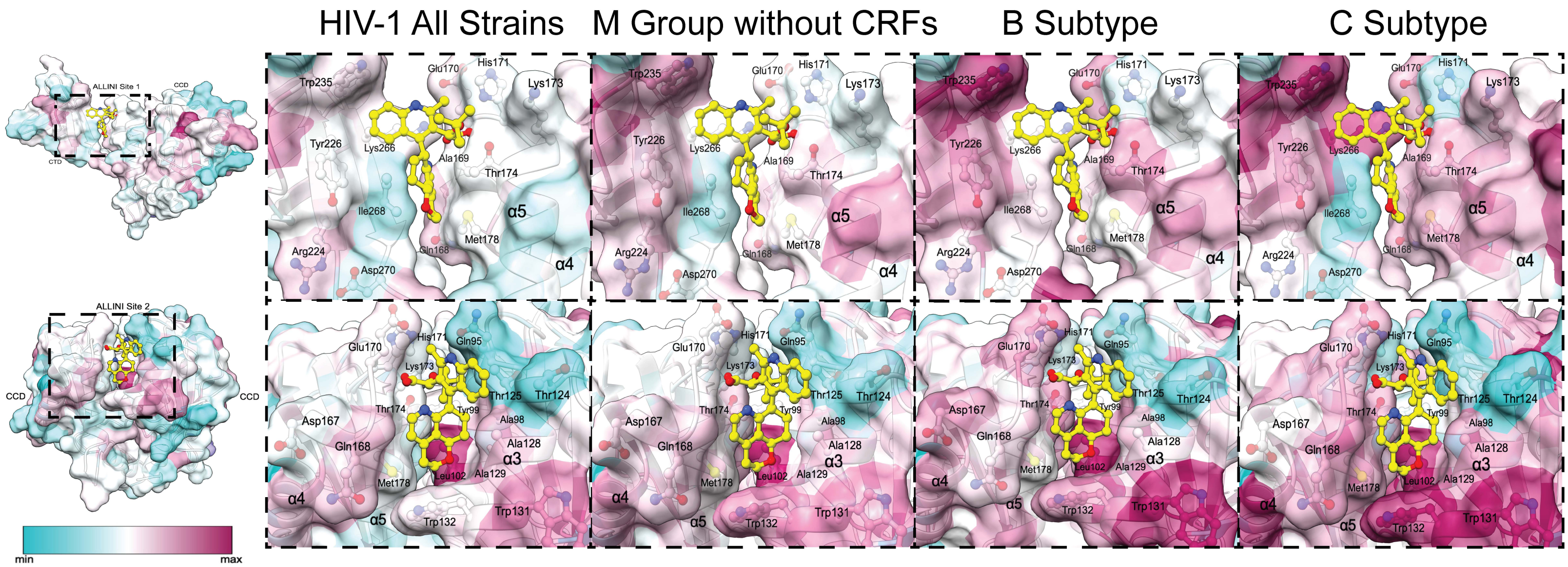


Supplemental Figure 3.
